# Rising out of the ashes: additive genetic variation for susceptibility to *Hymenoscyphus fraxineus* in *Fraxinus excelsior*

**DOI:** 10.1101/031393

**Authors:** F. Muñoz, B. Marçais, J. Dufour, A. Dowkiw

## Abstract

Since the early 1990s, ash dieback due to the invasive ascomycete *Hymenoscyphus fraxineus* is threatening *Fraxinus excelsior* in most of its natural range. Previous studies reported significant levels of genetic variability for susceptibility in *F. excelsior* either in field or inoculation experiments. The present study was based on a field experiment planted in 1995, fifteen years before onset of the disease. Crown and collar status were monitored on 788 trees from 23 open-pollinated progenies originating from 3 French provenances. Susceptibility was modeled using a Bayesian approach where spatio-temporal effects were explicitly taken into account. Moderate narrow-sense heritability was found for Crown Dieback (CD, h^2^=0.42). This study is first to show that Collar Lesions are also heritable (h^2^=0.49 for prevalence and h^2^=0.42 for severity) and that there is significant genetic correlation (r=0.40) between the severities of both symptoms. There was no evidence for differences between Provenances. Family effects were detected, but computing Individual Breeding Values (IBV) showed that most of the genetic variation lies within families. In agreement with previous reports, early flushing correlates with better crown status. Consequences of these results in terms of management and breeding are discussed.

## Introduction

An extensive review on the European ash dieback crisis was published recently (McKinney et al. 2014), thus allowing for a brief overview here. Severe dieback of European common ash (*Fraxinus excelsior*) was first reported in Poland and Lithuania in the early 1990s (Lygis et al. 2005; Przybyl 2002). The observed symptoms have long been thought to result from a combination of climatic factors and new pathogens and vectors (Pliūra and Baliuckas 2007). It was fourteen years after the first reports that an ascomycete was finally identified as the primary causal agent (Kowalski 2006). It is now present in at least 26 countries with its current South Western limit being Central France. Firstly described as a new fungal species (*i.e. Chalara fraxinea*), it was soon suggested that it could be the anamorphic stage of *Hymenoscyphus albidus*, a widespread native decomposer of ash litter (Kowalski and Holdenrieder 2009). However, further investigations concluded on a distinct and previously undescribed species now referred to as *H. fraxineus* (Baral et al. 2014). Recent findings suggest that the species is invasive and originates from Asia. First, the fungus could not be found in disease-free areas like Western and Central France (Husson et al. 2011) whereas it completely replaced *H. albidus* in infected areas of Denmark (McKinney et al. 2012b). Second, indications of a founder effect were reported at some loci in European samples (Bengtsson et al. 2012). Third, some fungi isolated from the Asian Ash species *Fraxinus mandshurica* in Japan and initially reported as *Lambertella albida* were subsequently identified as *H. fraxineus* (Zhao et al. 2012).

Knowledge on the life cycle of *H. fraxineus* has been improved and summarized by Gross *et al.* (2012). Sexual reproduction is hypothesized to be mediated through conidia which are produced in autumn on dead ash petioles in the litter and possibly on other dead tissues also. A single petiole can host multiple genotypes which overwinter in the form of black pseudosclerotial plates on which apothecia emerge in summer. Ascospores are wind-dispersed and germinate on ash leaflets or petioles forming an appressorium (Cleary et al. 2013). Once in the leaf tissue, the mycelium develops intracellularly, moving through the cells and easily colonizing the phloem, paratracheal parenchyma and parenchymatic rays (Dal Maso et al. 2012). Symptoms of the disease are many: wilting of leaves, necrotic lesions on leaves, necrosis on twigs, stems and branches, and collar lesions. Collar lesions have been described quite late (Lygis et al. 2005) and the question of their primary cause - *H. fraxineus* or other fungi like *Armillaria sp.* or *Phytophthora sp.* – is still controversial (Bakys et al. 2011; Enderle et al. 2013; Husson et al. 2011; Orlikowski et al. 2011; Skovsgaard et al. 2010).

With its natural range stretching from Iran to Ireland and from Southern Scandinavia to Northern Spain (Dobrowolska et al. 2011), *F. excelsior* is the most common and the most northern of the 3 *Fraxinus* species native to Europe, and thus the first one to face this new threat. Detailed quantifications are scarce, but consequences of the disease can be severe. In Lithuania, 10 years after the first report, over 30,000 ha of common ash stands were reported to be affected and mortality was estimated to be 60% state wide while in some parts of the country only 2% of the trees remained visually healthy (Juodvalkis and Vasiliauskas 2002). In 2006, ash dieback was still present on 9,400 ha despite extensive clear cuts of severely damaged stands (Pliūra and Baliuckas 2007). In Southern Sweden, one fourth of all ash trees were reported as dead or severely damaged 7 years after the first report (Fischer et al. 2010). In some parts of North-Eastern France, only 3% of the trees remained completely healthy 2 years after the first reports (Husson et al. 2012). These figures corroborate the findings from Mc Kinney *et al.* (2011) on 39 clones from 14 Danish populations where only one clone out of 39 maintained an average damage level below 10% in two replicated field trials. The computed broad-sense heritability estimates (*i.e.* 0.40-0.49) were however indicative of a strong genetic control, meaning that there is an adaptive potential and that breeding material with low susceptibility would be possible. Evidences of a significant level of genetic variation were also found in a few other comparison experiments involving clones or half-sib families (Pliūra et al. 2011; Kjær et al. 2012; Stener 2013; Pliūra et al. 2014; Lobo et al. 2015).

The present study explores the genetic variability of common ash for susceptibility to *H. fraxineus* using open-pollinated maternal progenies. It is based on a 20-year old field trial with 23 half-sib families from 3 French provenances. The trial is located in the area where Ash dieback was first detected in France in 2008. Because the stand has been monitored every year since disease appearance, spatio-temporal components of disease spread could be taken into account to help avoiding confusion between disease escape and resistance. Moreover, a Bayesian approach was used instead of the classical frequentist analysis. Bayesian methods accommodate complex spatio-temporal structures in a more straightforward way and their results are directly interpretable in terms of posterior probabilities. In addition, they are also more suitable for modeling data which deviates heavily from normality, as it is frequently the case during early stages of disease spread. In addition, it leverages information from previous knowledge with actual observed data, which can be critical in situations where data are little informative (Mila and Carriquiry 2004). This study is also first to report on the genetic parameters for collar lesions and on the genetic correlation with crown defoliation.

## Materials and Methods

### Experimental design

The studied material consisted in 23 open-pollinated *F. excelsior* maternal progenies from 3 North-Eastern French provenances (Amancey, 8 families; Chalèze, 7 families; Vernois-sur-Mance, 8 families). Although located less than 40 km apart (Figure 1), these provenances differ for elevation and soil composition and structure. Mother trees were selected at random within each provenance. Seeds were collected in 1992, cold-stratified, germinated and raised in nursery until plantation in January 1995.

**Figure 1.**
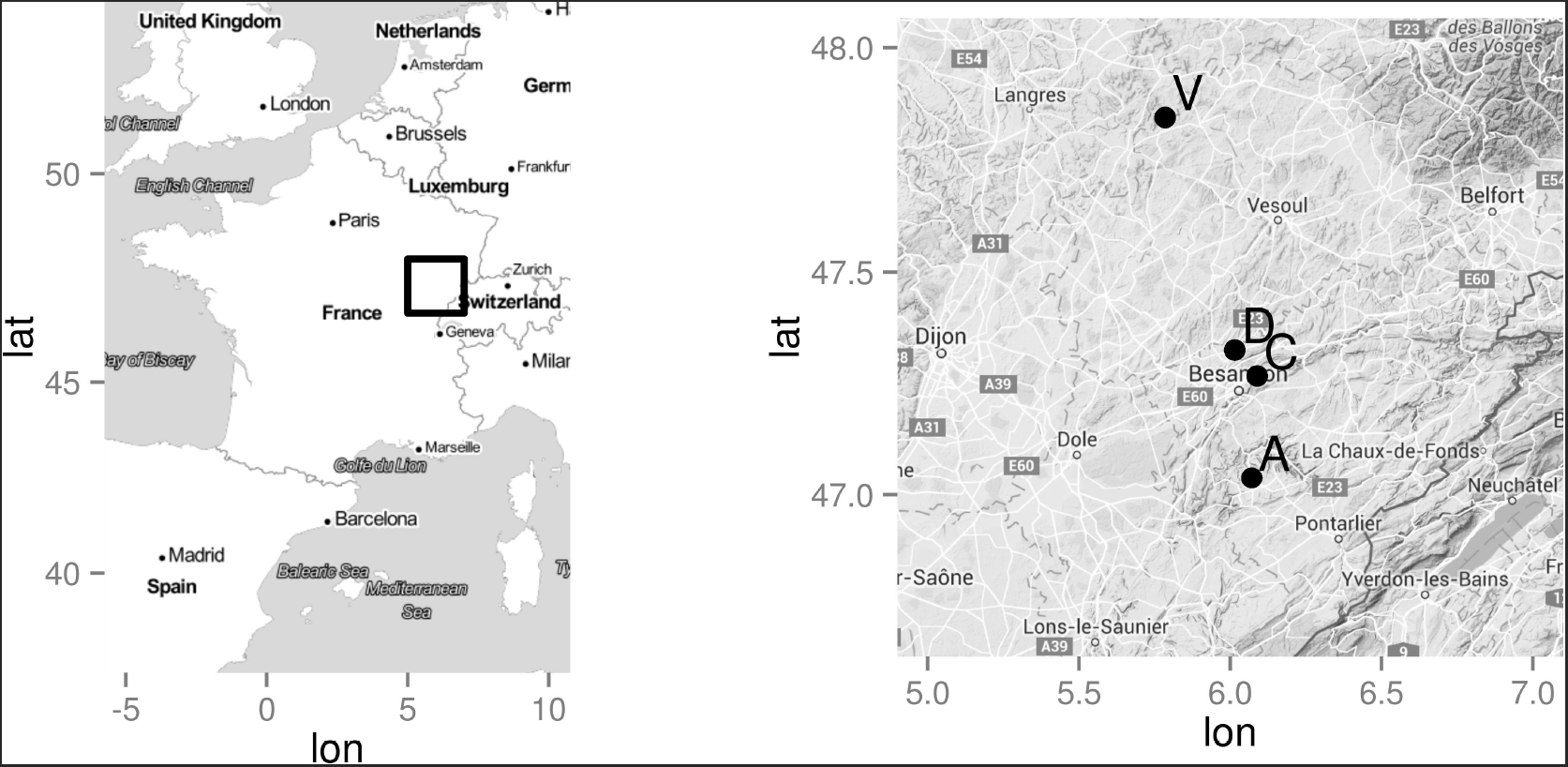
Provenance (A=Amancey, C=Chalèze, V=Vernois-sur-Amance) and study site (D=Devecey) locations.

The trial was established in Devecey, 61 km from the furthest provenance (Figure 1), on a winter tilled agricultural land located 250 m high. Planting was done at a spacing of 4 m x 4 m following a randomized incomplete block design. Depending on plant availability, each family was represented in 2 to 10 blocks plus the border except two families that were represented in the border only. Each block contained 16 families, each represented by 4 trees that were distributed among 4 sub-blocks. Each family was thus represented by 8 to 68 trees in the whole trial including the border (Table 1). nostar

**Table 1:** Studied *F. excelsior* material

| Provenance | Half-sib family<sup>a</sup> | Number of blocks of the family is represented in<sup>b</sup> | Total number of trees |
| --- | --- | --- | --- |
| Amancey | (A03) | 2 + | 11 |
|  | (A04) | 0 + | 8 |
|  | A09 | 10 + | 49 |
|  | A10 | 8 + | 40 |
|  | A12 | 7 + | 30 |
|  | A14 | 7 + | 29 |
|  | A21 | 10 + | 60 |
|  | A23 | 7 + | 30 |
| Chalèze | C01 | 10 + | 48 |
|  | C03 | 10 + | 65 |
|  | C08 | 10 + | 68 |
|  | C09 | 9 + | 37 |
|  | C13 | 10 + | 49 |
|  | C16 | 10 + | 41 |
|  | C22 | 5 + | 21 |
| Vernois-sur-Mance | V01 | 5 + | 24 |
|  | (V05) | 0 + | 10 |
|  | (V10) | 3 | 12 |
|  | V17 | 10 + | 51 |
|  | (V18) | 3 | 12 |
|  | V21 | 5 + | 24 |
|  | (V22) | 4 | 16 |
|  | V23 | 10 + | 49 |
| Total |  |  | 788<sup>c</sup> |

<sup>a</sup> Families in parenthesis were represented in less than 5 blocks
<sup>b</sup> The + symbol indicates that the family is represented in the border of the experiment
<sup>c</sup> Four trees for which provenance information was lost are located in the border and included in this number

### Measurements

A first investigation for the presence of *H. fraxineus* was conducted in February 2009 and did not reveal any symptom. The first signs of infection were observed in February 2010 and presence of the fungus was confirmed by real time PCR (French Department of Agriculture, personal communication). Crown dieback (CD) was measured in July 2010, July 2011, June 2012, June 2013 and June 2014 using a 0 to 5 ranking scale according to the proportion of dead branches: 0 - no dead branches; 1 - less than 10% dead branches; 2 - 10 to 50% dead branches; 3 - 50 to 80% dead branches; 4 - more than 80% dead branches; 5 - dead tree. For data analysis, classes were subsequently converted to their median value (i.e. 0, 0.05, 0.30, 0.65, 0.90 and 1). Collar lesions (CL) were measured in July 2012 and June 2013. Detecting and measuring them required scratching the bark using a triangular scraper; this is why this measurement was conducted for 2 years only. As several lesions can occur on the same tree, basal width of all lesions were cumulated and divided by the basal girth (BG) of the tree to compute CL as a 0 to 1 girdling index (Figure 2). Bud flushing was measured once, three years after planting, using a 1 (late) to 5 (early) ranking scale.

**Figure 2:**
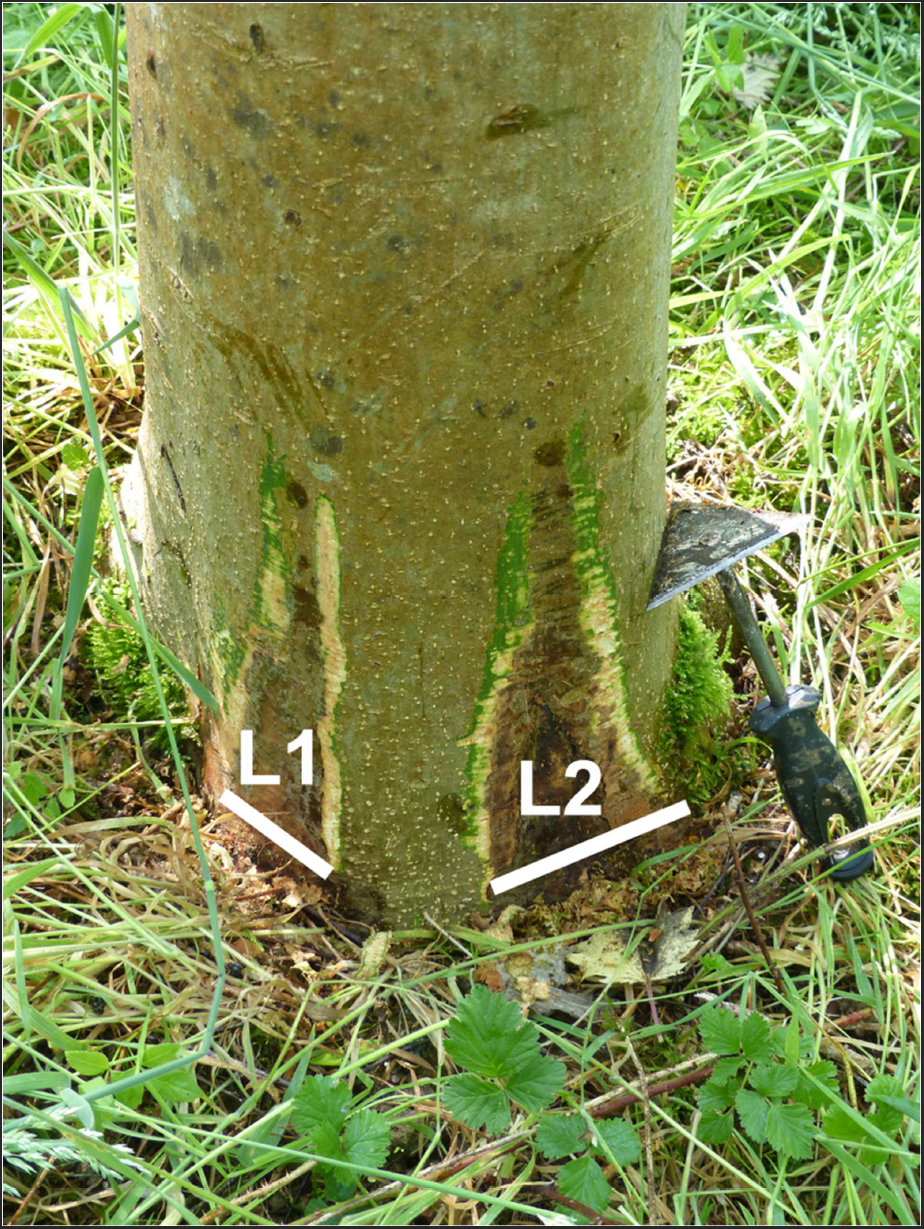
Collar lesions (CL) were cumulated and expressed as a girdling index (i.e. *L*_1_ + *L*_2_ + … + *L*_*n*_ over the basal circumference of the tree).

### Data analysis

We analyzed each symptom independently using a sequence of statistical models of increasing complexity and flexibility. All of the fitted models for CD belong to the family of Linear Mixed Models (LMM). This is, the vector *y* of *n* measurements of phenotypic values are described as a linear model with a vector of *p* fixed effects (*β*) and a vector of *q* random effects (*u*). In matrix form,

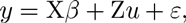

where X and Z are *n × p* and *n × q* incidence matrices, respectively, and *ε* is the vector of residuals. Furthermore, both *u* and ε are assumed to be independent from each other and to follow a zero-mean Gaussian distribution with covariance matrices 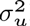R and σ^2^I respectively, where R is a *q × q* structure matrix, and I is the *n × n* identity matrix.

Several independent random effects with specific variances and structure matrices can be stacked in a single vector *u* with a block-diagonal covariance matrix where each block is given by the corresponding component and an incidence matrix *Z* binding the individual incidence matrices column-wise.

The normality of the residuals is a delicate assumption to make for response variables that are categorical (CD) or restricted to the interval 0-1 (CL). However, it is very convenient from a computational point of view, and this approximation is commonly found in other studies (Kjær et al. 2012; Pliūra et al. 2011). We performed a transformation of the CD variable to improve the adjustment to this hypothesis. Specifically, we worked with

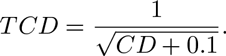

No normalizing transformation was found for CL. In consequence, we used a mixture of Generalized Linear Mixed Models (GLMM) for this trait. A GLMM is an extension of a LMM for non-Gaussian data. The observations are assumed to be an independent random sample from some distribution of the exponential family conditional to the mean, which is modeled as a non-linear function of the linear predictor:

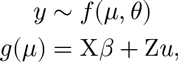

where *g* is an appropriate link function, *μ* is the mean, and *θ* is a vector of additional distribution-specific parameters.

### Model inference and comparison

All analyses were performed using R (R Core Team 2015). The models were fitted from a Bayesian perspective using the Integrated Nested Laplace Approximation (INLA) methodology (Rue and Martino 2009) and software (Rue et al. 2014). The Marginal Likelihood, the DIC (Spiegelhalter et al. 2002) and the WAIC (Watanabe 2010) were used as model selection criteria. The Marginal Likelihood was scaled by a factor of –2 (also known as the Deviance), so that for all three criteria, lower values are better.

### Statistical models for Crown Dieback

An appropriate statistical model was determined following a model-selection procedure (Table 3 in the Annex). The initial reference model included only unstructured random effects (*i.e.* the matrix R is an identity matrix for all random effects) and is analogous to what has been commonly used in previous studies on the genetic diversity of resistance to *H. fraxineus* (Kjær et al. 2012; Pliūra et al. 2011). The data was best described by a model including fixed effects for the Year and Bud-Flush precocity, a random additive genetic effect at individual level and a spatio-temporal (ST) random effect (see Table 3 in the Annex and details therein).

**Table 3:** Model comparison for TCD. The variables in the *Linear Predictor* are the year, the block, the family (Fam), the family by block interaction, the individual breeding value (IBV), the spatio-temporal effect (ST), the provenance (Prov) and the bud flush precocity (BF).

| Model | Specificity | Linear predictor<sup>a</sup> | DIC | p_D | WAIC | p_W | Dev. |
| --- | --- | --- | --- | --- | --- | --- | --- |
| M1 | Pliura et al. 2011 | Year + Block + *Fam* + *Fam:Block* | 5768 | 132 | 5766 | 126 | 6113 |
| M2 | Individual genetic effect | Year + Block + *IBV* | 4548 | 619 | 4573 | 568 | 5589 |
| M3 | Spatio-temporal effect | Year + *ST* + *IBV* | 4277 | 749 | 4307 | 666 | 5286 |
| M4 | Fixed provenance effect | Year + Prov + *ST* + *IBV* | 4278 | 732 | 4310 | 661 | 5314 |
| M5<sup>b</sup> | Fixed budflush effect | Year + BF + *ST* + *IBV* | 4272 | 745 | 4304 | 671 | 5325 |
| M6 | Null variance between families | Year + BF + *ST* + (*Fam=0*) + *IBV* | 3858<sup>c</sup> | 1706 | 3851<sup>c</sup> | 1326 | 5633 |

DIC: Deviance Information Criterion; p\_D: effective number of parameters for DIC; WAIC: Watanabe-Akaike information criterion; p\_W: effective number of parameters for WAIC; Dev.: Deviance = -2\*marginal log-likelihood.
<sup>a</sup> Fixed effects are represented in straight font, while random effects are shown in italics.
<sup>b</sup> Selected model is M5.
<sup>c</sup> The values of DIC and WAIC for model M6 are not reliable.

The additive-genetic individual effect is a structured random effect with a known covariance structure given by the family kinship. Specifically, the covariance matrix is

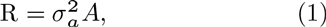

where 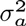 is the unknown additive genetic variance in the base population and the additive-genetic structure matrix A has elements *A_i j_* = 2Θ*_i j_*, i.e., twice the coefficient of coancestry between the individuals *i* and *j* (see for example Lynch and Walsh 1998).

The Spatio-Temporal random effect (ST) was modeled based on a Gaussian spatial process evolving in time, thus allowing for continuous environmental variation. Evaluated in the spatiotemporal locations of the observations, the values of the Gaussian process follow a multivariate Normal distribution with a covariance structure given by the distance, in space and time, between observations. Specifically, the spatio-temporal structure is built as the Kronecker product of separate temporal and spatial processes. The temporal process is simply determined by an autocorrelation parameter *ρ_t_* between observations, while the spatial process is defined by a Matérn covariance function with parameters of shape (*i.e*, smoothness) *v*, spatial scale *κ* and precision *τ*^2^. The smoothness parameter was fixed to *v* = 1 for convenience. The spatial scale parameter is associated with the effective range of the spatial process, so that the correlation between locations at a distance 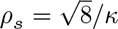 is approximately 0.13. Finally, the marginal variance of the spatial process is given by 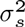 = 1/(4*πκ*^2^*τ*^2^). This yields a structured random effect with a parametric covariance matrix as follows:

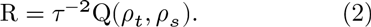

Consequently, while the global temporal trend is captured by the explicit Year effect in this model, the ST structure accounts for heterogeneous spatial deviations from the main trend both in space and time.

Furthermore, we also fitted a re-parameterization of this model introducing an explicit Family effect allowing to split the genetic variance into the *interfamily* and intra-family components and to compare their relative magnitudes. Computationally, this requires introducing sum-to-zero constraints for each family in the additive-genetic effect for preserving the identifiability of the model.

### Statistical models for Collar Lesions

This variable was considered as the result of two different processes or stages. First is the process determining whether a tree becomes infected or not. Second is the process determining how severely an infected tree is affected by the disease. Different factors, including the genetics and the microenvironment, could potentially affect the two processes in different ways. Therefore, the two processes were analyzed separately.

The pattern of zeros (*i.e.* inversely related with the disease prevalence) was modeled with a Bernoulli likelihood while the strictly positive observations (*i.e.* disease severity) were assumed to follow a continuous distribution with positive support. For the continuous component, Beta and Gamma were considered as candidate distributions and a preliminary assessment was performed to determine which one fitted the data better. All the potentially explanatory fixed effects were included for this preliminary analysis (Table 5 in the Annex). The Gamma distribution emerged as the best candidate and was used in all subsequent models.

All other models systematically included an individual additive-genetic and a Spatio-Temporal (ST) effect as defined in the previous subsection. Two separate variable selection procedures were conducted for the binary and continuous components of CL to assess the relevance of the Basal Circumference (BC) and the Provenance (Tables 4 and 5 in the Annex, respectively).

**Table 4:** Model comparison for the binary component of CL. The variables in the *Linear predictor* are the year, the spatio-temporal effect (ST), the individual breeding value (IBV), the Basal Circumference (BC) and the provenance (PROV).

| Model | Specificity | Linear predictor<sup>a</sup> | DIC | p_D | WAIC | p_W | Dev. |
| --- | --- | --- | --- | --- | --- | --- | --- |
| M0<sup>b</sup> | Reference | Year + *ST* + *IBV* | 1498 | 356 | 1443 | 249 | 1978 |
| M1.1 | BC categories | Year + BC + *ST* + *IBV* | 1485 | 368 | 1428 | 255 | 2014 |
| M1.2 | BC cat. x Year | Year * BC + *ST* + *IBV* | 1472 | 386 | 1410 | 264 | 2051 |
| M1.3 | BC linear | Year + $\beta$ BC + *ST* + *IBV* | 1494 | 361 | 1438 | 251 | 1988 |
| M1.4 | BC quadratic | Year + $\beta_0$ BC + $\beta_1$ BC<sup>2</sup> + *ST* + *IBV* | 1492 | 361 | 1436 | 251 | 1996 |
| M1.5 | BC non-param. | Year + f(BC) + *ST* + *IBV* | 1494 | 360 | 1438 | 251 | 2254 |
| M2.1 | PROV | Year + PROV + *ST* + *IBV* | 1493 | 366 | 1434 | 253 | 1994 |
| M2.2 | PROV x Year | Year * PROV + *ST* + *IBV* | 1498 | 367 | 1441 | 254 | 2013 |

DIC: Deviance Information Criterion; p\_D: effective number of parameters for DIC; WAIC: Watanabe-Akaike information criterion; p\_W: effective number of parameters for WAIC; Dev.: Deviance = -2\*marginal log-likelihood. $\beta$ , $\beta_0$ and $\beta_1$ are regression coefficients, and f(·) represents an unknown smooth function.
<sup>a</sup> Fixed effects are represented in straight font, while random effects are shown in italics.
<sup>b</sup> Selected model is M0.

**Table 5:** Model comparison for the continuous component of CL. The variables in the *Linear predictor* are the family (FAM), the provenance (PROV), the year, the Basal Circumference (BC), the spatio-temporal effect (ST) and the individual breeding value (IBV).

| Model | Specificity | Linear predictor<sup>a</sup> | DIC | p_D | WAIC | p_W | Dev. |
| --- | --- | --- | --- | --- | --- | --- | --- |
| M0.1 | Beta likelihood | FAM + PROV + Year * BC | -600 | 34 | -605 | 27 | -302 |
| M0.2 | Gamma likelihood | FAM + PROV + Year * BC | -763 | 34 | -757 | 37 | -459 |
| M0.3<sup>b</sup> | Reference | Year + *ST* + *IBV* | -1144 | 289 | -1138 | 232 | -661 |
| M1.1 | BC categories | Year + BC + *ST* + *IBV* | -1140 | 188 | -1133 | 231 | -616 |
| M1.2 | BC cat. x Year | Year * BC + *ST* + *IBV* | -1136 | 193 | -1131 | 234 | -573 |
| M1.3 | BC linear | Year + $\beta$ BC + *ST* + *IBV* | -1140 | 187 | -1133 | 231 | -650 |
| M1.4 | BC quadratic | Year + $\beta_0$ BC + $\beta_1$ BC<sup>2</sup> + *ST* + *IBV* | -1140 | 188 | -1134 | 231 | -638 |
| M1.5 | BC non-param. | Year + f(BC) + *ST* + *IBV* | -1143 | 190 | -1136 | 233 | -379 |
| M2.1 | PROV | Year + PROV + *ST* + *IBV* | -1094 | 316 | -1112 | 234 | -614 |
| M2.2 | PROV x Year | Year * PROV + *ST* + *IBV* | -1115 | 286 | -1107 | 231 | -601 |

DIC: Deviance Information Criterion; p\_D: effective number of parameters for DIC; WAIC: Watanabe-Akaike information criterion; p\_W: effective number of parameters for WAIC; Dev.: Deviance = -2\*marginal log-likelihood. $\beta$ , $\beta_0$ and $\beta_1$ are regression coefficients, and f(·) represents an unknown smooth function.
<sup>a</sup> Fixed effects are represented in straight font, while random effects are shown in italics.
<sup>b</sup> Selected model is M0.3.

Finally, both components of CL were integrated into a final model which consisted into a mixture of both GLMMs. For a measurement *y_i j k_* taken in year *i*, at the location *j* for the individual *k*, we assumed that

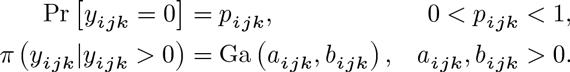

We specified a hierarchical model for the parameters *p_ijk_*, *a_ijk_* and *b_ijk_* using appropriate link functions of the expected values of the respective distributions. Specifically, calling *μ* = E [*y*|*y* > 0] = *a/b*, we defined two linear predictors

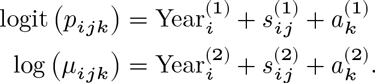

For each linear predictor, Year¿ is the fixed effect of the year *i* = 2012, 2013; *s_ij_* is a structured Spatio-Temporal (ST) random effect and *a_k_* is a structured additive-genetic random effect at individual level (*i.e.* a vector of Individual Breeding Values, IBV). The ST structure was built as the Kronecker product of separate temporal and spatial zero-mean Gaussian processes, as in Eq. 2. Finally, the structured additive-genetic effect followed a zero-mean multivariate Normal distribution with a known covariance structure given by the family kinship, as in Eq. 1.

### Prior distributions

All the fixed effects had a vague zero-mean Gaussian prior with variance of 1000.

For the variance 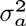 of the additive-genetic effect we used an inverse-Gamma with shape and scale parameters of 0.5. This is equivalent to an Inverse-Chi-Square with 1 df, and places the 80% of the density mass between 0.05 and 15, with a preference for lower values.

For the ST structure, the priors were set independently for the spatial and temporal structures. INLA provides a bivariate Gaussian prior for the logarithm of the positive parameters *κ* and *τ*^2^ of the spatial Matérn field. Its mean and variance were chosen to match reasonable prior judgments about the range and the variance of the spatial field. Specifically, the spatial range was concentrated between 5 and 38 tree spacings, the latter being the length of the shortest side of the field. There might well exist very short-ranged environmental factors affecting one or two trees at a time, but the model would not be able to separate them from random noise. On the other extreme, there certainly are environmental factors with a larger range than the field dimensions, but again, this would be virtually indistinguishable from the global mean of the field. Moreover, the results were found to be very robust to small variations of these prior statements.

### Heritability estimates

In general, the narrow-sense heritability is computed as the ratio between the additive-genetic variance and the phenotypic variance. However, this can be implemented in practice in different ways, depending on the specific model and the researcher goals and criteria.

The selected models for CD and for both components of CL yield direct estimates of the additive-genetic variance.

For CD, the phenotypic variance is computed as the sum of the variance components of the model. Namely, the additive-genetic variance, the residual variance, and following accepted guidelines (Visscher et al. 2008), the variance of the ST effect.

Estimating the heritability of the CL symptom is more complex. First, because there are two different traits in the model, and thus two different measures of heritability. But more importantly, because both with Binomial and Gamma likelihoods, the phenotypic variance is a function of the mean, and thus varies from observation to observation. Furthermore, although the additive-genetic variances are assumed common to the population, the phenotypic variance cannot be decomposed additively into genetic and residual components. Most approaches in the literature dealing with binary variables use the so-called *Threshold Models*, where the data are assumed to be deterministically 0 or 1 depending on whether some unobserved latent Gaussian variable reaches some threshold (see, for example, Dempster and Lerner 1950). In our case the response variable given the latent structure is random rather than deterministic. Therefore, the residual variability comes from the likelihood distribution, which is in a different scale than the genetic variance parameter. However, this is equivalent to a threshold model with an additional residual term following a Logistic distribution. This results in the following formula for the heritability of the binomial component in the latent scale:

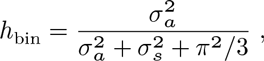

where *π*^2^/3 is the variance of a Logistic distribution (Nakagawa and Schielzeth 2010), and 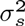 is the variance of the ST effect. For the continuous component we followed the general simulation-based approach for a GLMM described in de Villemereuil et al. (2015).

All heritability estimates have been computed by Monte Carlo simulation given the posterior distributions of the relevant variance parameters. Specifically, we sampled 5000 independent observations of each variance from their posterior distribution, and derived a posterior density for the her-itability.

## Results

### Crown dieback occurrence and severity

From 2010 to 2014, mean CD increased from 0.01 to 0.27. The distribution of individual values remained highly skewed to the right during the whole study time with most trees showing less than 50% dead branches (i.e. CD<0.5, Figure 3). In 2010, when disease started to be seen, 53 trees (i.e. 7%) had visible crown dieback including four trees with CD> 0.5. Although one of these four trees died in less than one year, the proportion of dead trees reached only 3% in 2014 while the proportion of trees with CD > 0.5 increased to 19%. The annual decrease in the proportion of trees without visible CD was similar in 2011 and 2012 (-24% and -26%, respectively) but it accelerated in 2013 (-53%) and again in 2014 (-74%), leading to a situation where only 49 trees (i.e. 6%) remained free from CD in 2014. When computing the 3104 values of individual annual increments for CD over the four years of the study, only 73 negative values were found (i.e. 2.3%).

**Figure 3:**
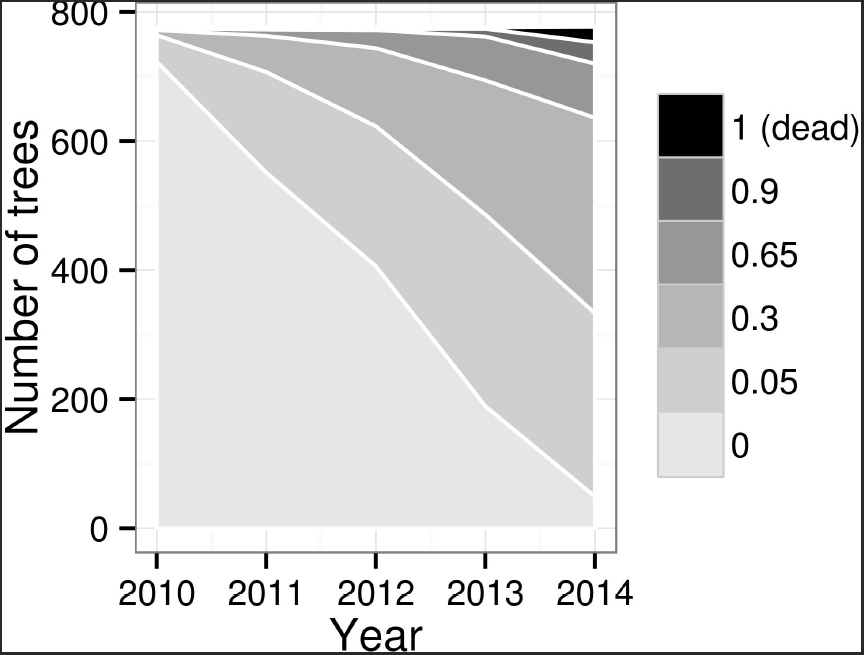
Crown dieback (CD) progress over time.

### Collar lesions occurrence and severity

From 2012 to 2013, mean CL increased from 0.05 to 0.14. Collar lesions were found on 29% of the trees in 2012 and this proportion doubled in just one year. Mean CL values computed on symptomatic trees only were 0.19 and 0.23 for 2012 and 2013, respectively with only 24 trees with CL > 2013 (Figure 4).

**Figure 4:**
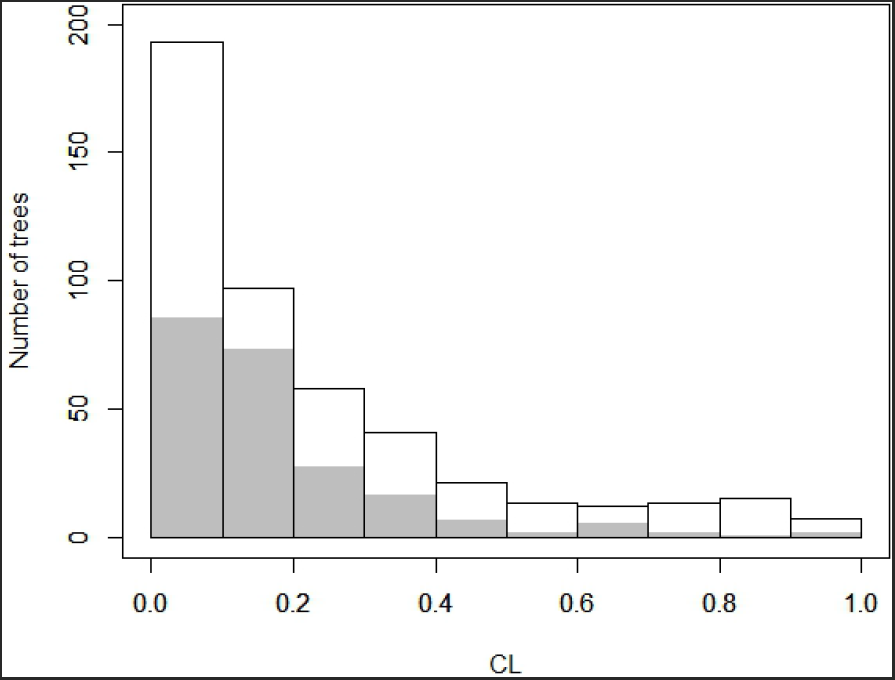
Distribution of individual values for collar lesions (CL) expressed as a 0-1 girdling index in 2012 (grey bars) and in 2013 (white bars). Individuals without collar lesions (CL=0) are not shown (*i.e.* 550 trees in 2012 and 305 trees in 2013).

Among them, four had their collar totally rotten in 2013 and they were dead the next year. Fifty eight negative values were found for the 2012-2013 increment in CL with only 12 increments below - 0.1 (i.e. 1.5 % of all computed increments). Most of these negative increments were probably artifactual due to the impossibility to measure basal girth and lesion length at exactly the same level each year due to the curved shape of the trunk basis.

Although a positive correlation was found between Basal Circumference (BC) and CL occurrence in 2012 (r_Spearman = 0.94, p-value = 0.017, Figure 5), collar lesion occurrence was not clearly related to basal circumference, considering the model comparisons conducted in Tables 4 and 5 in the Annex. In 2013, only the very few trees with basal circumference below 30 cm had significantly fewer basal cankers.

**Figure 5:**
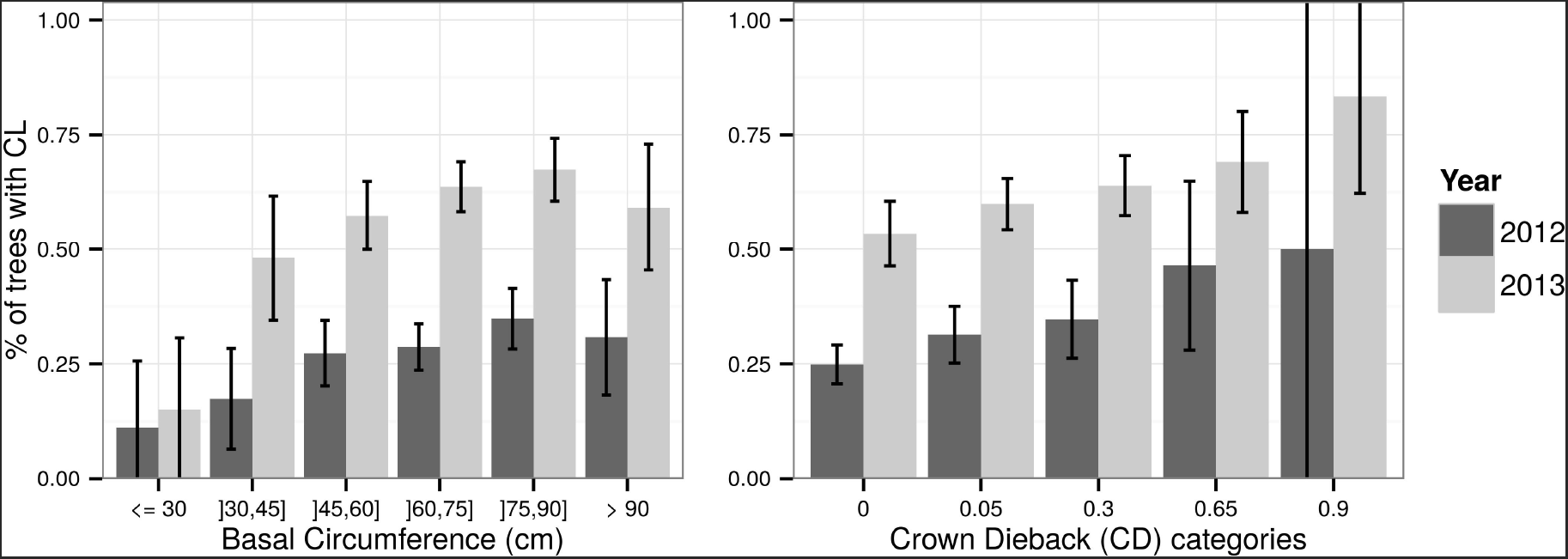
Occurrence of collar lesions (CL) according to basal circumference categories and Crown Dieback (CD) severity by year. Error bars = 95% confidence intervals for the proportions, considered independently from each other.

### Phenotypic correlation between symptoms

A positive correlation between occurrence of CL and CD severity was observed both in 2012 and 2013(r_Spearman = 1, p-value = 0.017; Figure 5). Despite this trend, collar lesions could be found in many trees without visible CD symptoms. Indeed, 53% of the trees with CD=0 in 2013 had collar lesions. Conversely, 71% of the trees with CL=0 in 2013 had crown dieback.

Severities of both traits were significantly correlated also, and correlation increased from 2012 to 2013 (Table 2)

**Table 2:** Phenotypic correlations between crown dieback (CD) and collar lesions (CL) symptoms. Unreported p-values correspond to values under the numerical precision.

|  | 2012 |  | 2013 |  | 2012 and 2013 |  |
| --- | --- | --- | --- | --- | --- | --- |
|  | Sample correlation | p-value | Sample correlation | p-value | Sample correlation | p-value |
| Pearson | 0.14 | 3.2e-05 | 0.31 | . | 0.29 | . |
| Spearman | 0.13 | 2.4e-04 | 0.21 | 4.0e-09 | 0.23 | . |
| Kendall | 0.11 | 2.6e-04 | 0.17 | 3.5e-09 | 0.20 | . |

Of the 22 trees that were dead in 2014 but still alive in 2013, 21 had CD values higher or equal to 0.65 in 2013 with fourteen of them having CL values higher than 0.5 in the same year. Interestingly, one of these 22 trees had no CL and a CD value of only 0.3 in 2013 but this was a small tree (basal circumference 34 cm).

### Modeling CD and estimating genetic parameters

Table 3 (in the Annex) summarizes the variable-selection procedure for TCD. The provenance was not found to be a relevant explanatory variable according to all model-comparison criteria. By contrast, there were clear differences in the mean genetic merit between families (Figure 6), although most of the genetic variability occurs within families (Figure fig:cd-variances-posteriors). Finally, although the relevance of bud-flush precocity was not conclusive in terms of the model-comparison criteria, prior evidence in the literature of a relationship between BF and CD (Bakys et al. 2013; McKinney et al. 2011; Pliūra and Baliuckas 2007; Stener 2013) supported its inclusion in the model. The fixed Year effect estimates revealed a clear progression of the disease, almost doubling the predicted CD value for a mean individual each year. Specific mean predicted values were 1.0% (2010); 2.6% (2011); 5.2% (2012); 10.7% (2013) and 17.8% (2014) with obvious variation according to Bud Flush precocity (BF, Figure 8). Early flushers (high BF values) tended to have lower CD values.

**Figure 6:**
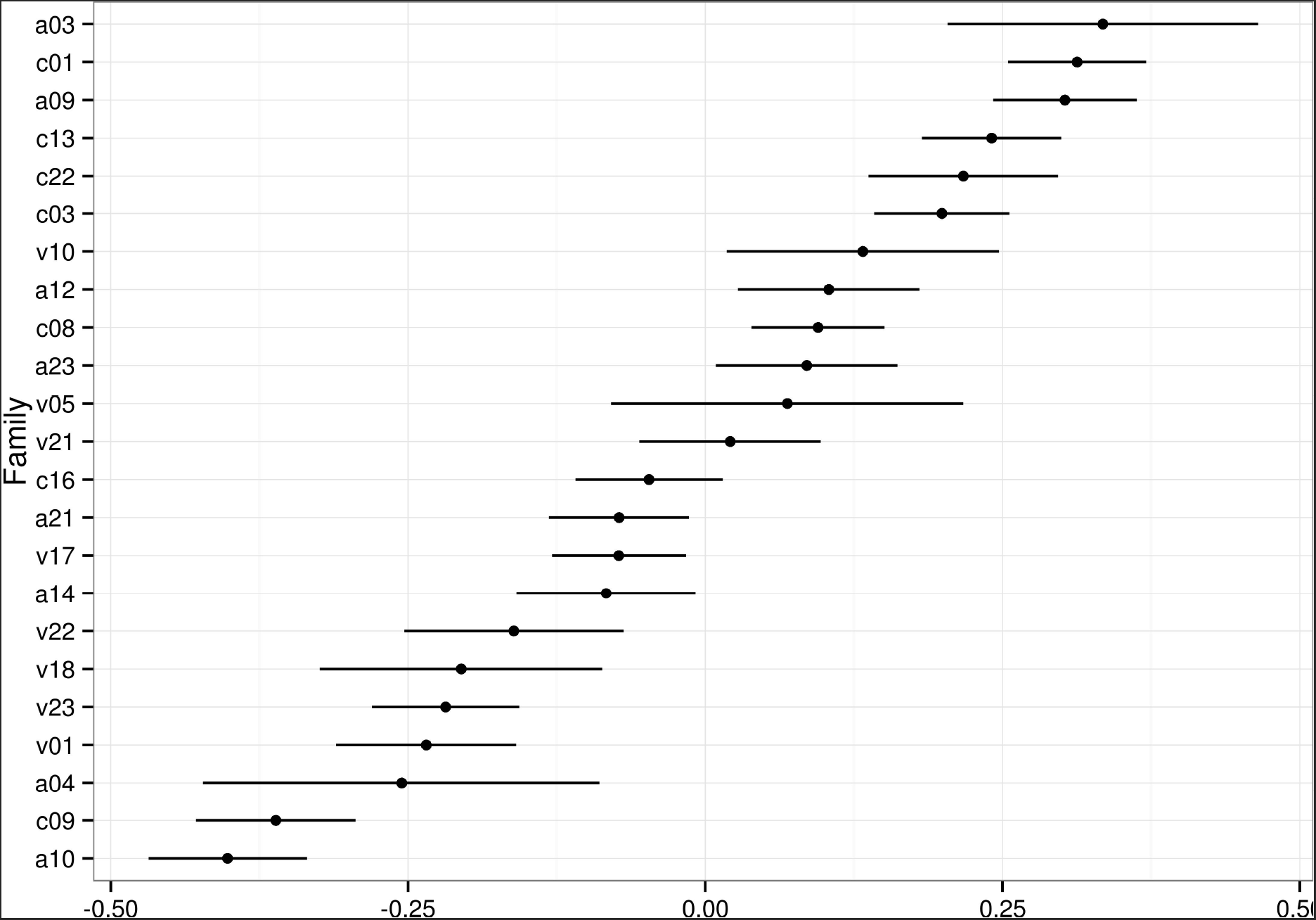
Posterior means and 95% Credible Intervals for the Family effects for Crown Dieback (transformed: TCD, families on the left are the most susceptible).

**Figure 8:**
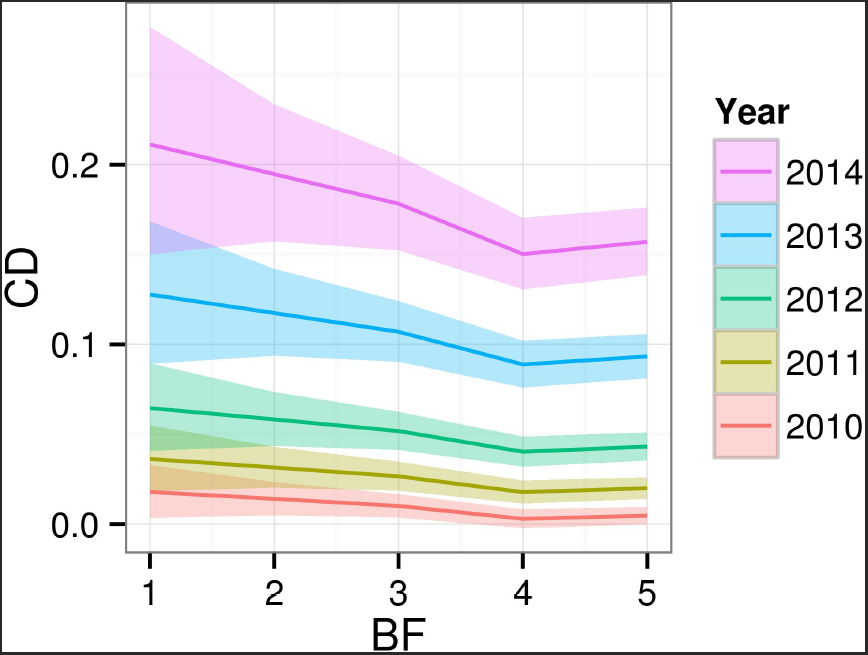
Posterior mean and 95% Credible Intervals effect of Bud Flush precocity (BF) on Crown Dieback (CD, original scale) by Year.

Figure 9 shows the posterior mean ST effect. The scale being inverted by the transformation (TCD), the disease in terms of CD appeared to be consistently more intense in the bottom-left corner of the field. The fact that the posterior distributions of the range and variance of the Gaussian process were concentrated well within the support of their corresponding prior indicated that the data were informative enough and the results were not constrained by the choice of the priors (Figure 12, in the Annex).

**Figure 9:**
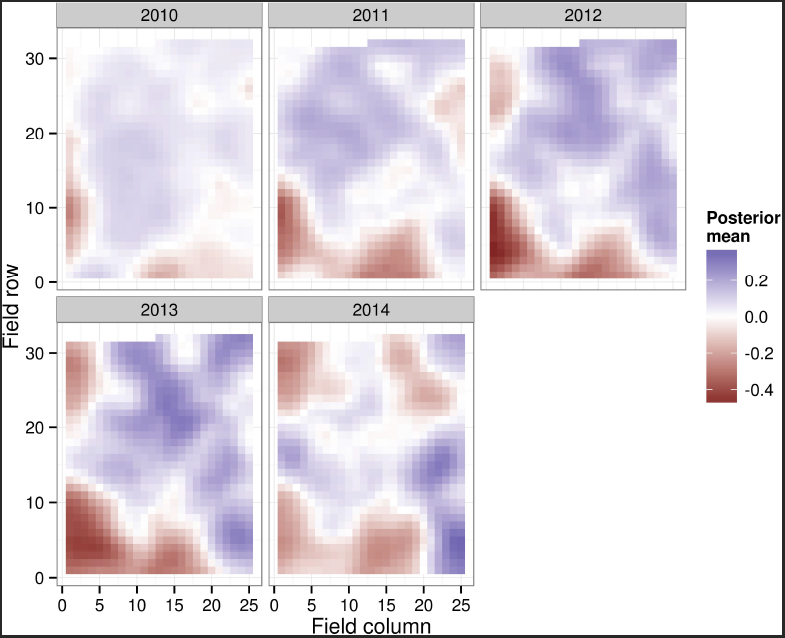
Posterior mean Spatio-Temporal (ST) gaussian fields for Crown Dieback (transformed: TCD, low values refer to high susceptibility).

**Figure 12:**
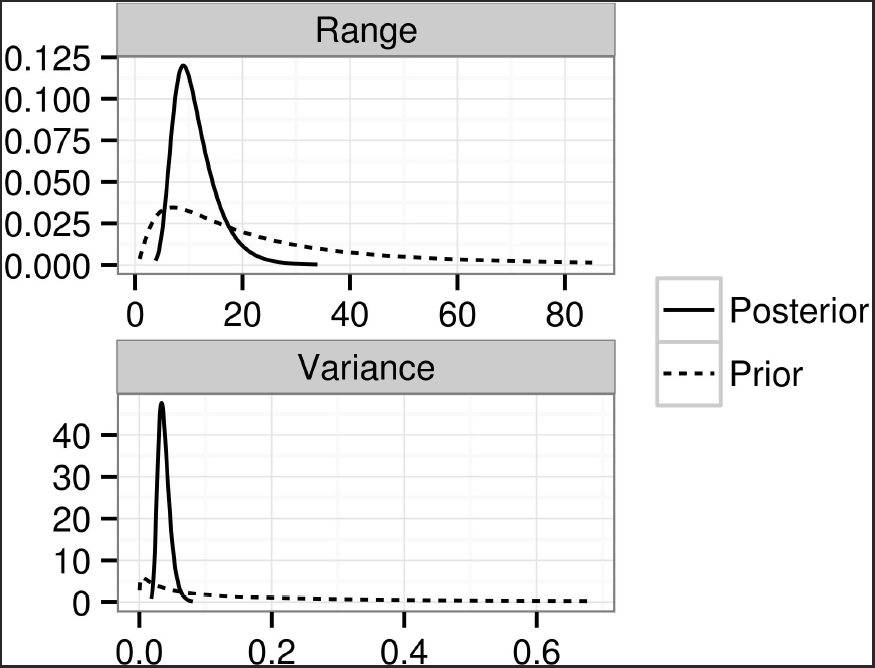
Prior and posterior densities for the spatial field characteristics for Crown Dieback (transformed: TCD). Range unit = spacing between trees.

The posterior modes and 95% HPD Credible Intervals were 0.137 (0.119, 0.156) for the additive-genetic variance and 0.147 (0.139, 0.157) for the residual variance. The estimates of the narrow-sense heritability *h^2^* were 0.42 (0.38, 0.47). Excluding the ST variance from the denominator increases the estimates up to 0.48 (0.44, 0.52).

### Modeling CL and estimating genetic parameters

Tables 4 and 5 (in the Annex) present the selection criteria for the competing models for the binary and continuous components of the full model for CL, respectively. For the binary component, including the Basal Circumference (BC) improved DIC and WAIC, particularly in categorical form with an interaction with the Year. However, it worsened the Deviance. Suspecting some confounding with the ST effect (see Discussion), this variable was not included in the final model. On the other hand, including a Provenance effect was clearly not relevant according to all three criteria. Neither BC nor the Provenance improved the model for the continuous component.

The plot of predicted against observed values for the binary and continuous components of the mixture model indicated a reasonable goodness of fit of the model (Figure 13, in the Annex). For the continuous component, the model fit yielded a prediction a bit shrinked, but this is expected due to the difficulty to predict values in the extremes of a bounded variable.

**Figure 13:**
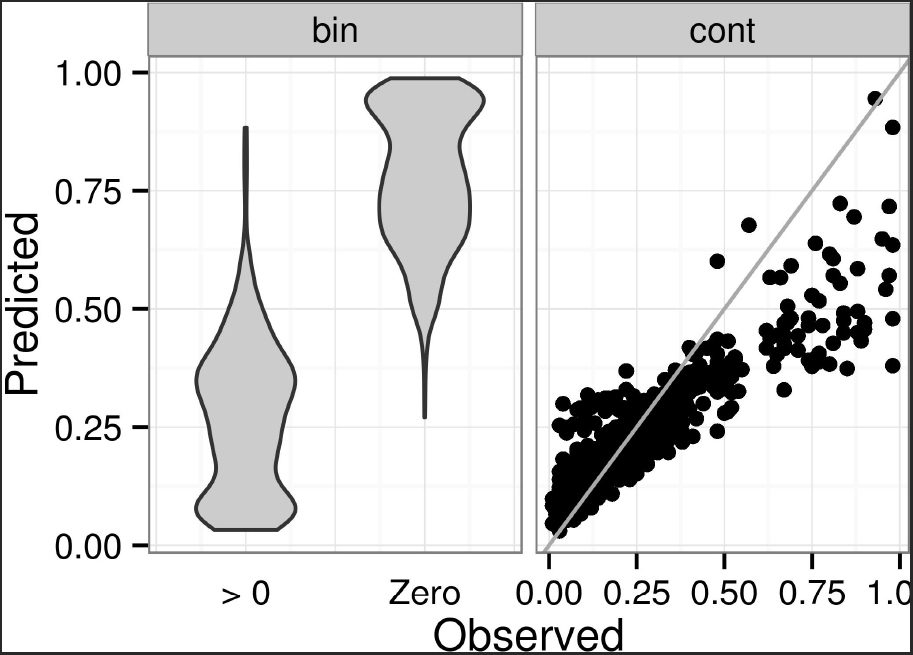
Fitted vs. observed values for the binomial (bin) and continuous (cont) components of the model for Collar Lesions (CL). In the binary component, the predicted value corresponds to the probability of an outcome of zero (i.e., complete absence of collar lesion).Since the binary observations clump together in only two values, we improved the visualization using a violin plot, a modification of a box plot which represents a density estimation of the data.

The distribution of mean posterior IBV for the binary component by infection status (classified in three categories) was as expected. Trees that did not show any sign of infection got the highest predicted values, and therefore an increased probability of an outcome of zero (Figure 14), while those that showed some level of infection both in 2012 and 2013 got the smallest IBV. The degree of overlapping between these histograms indicates the relative importance of the genetic component with respect to the rest, as a predictor of the phenotype. This is related with the heritability which is presented below.

**Figure 14:**
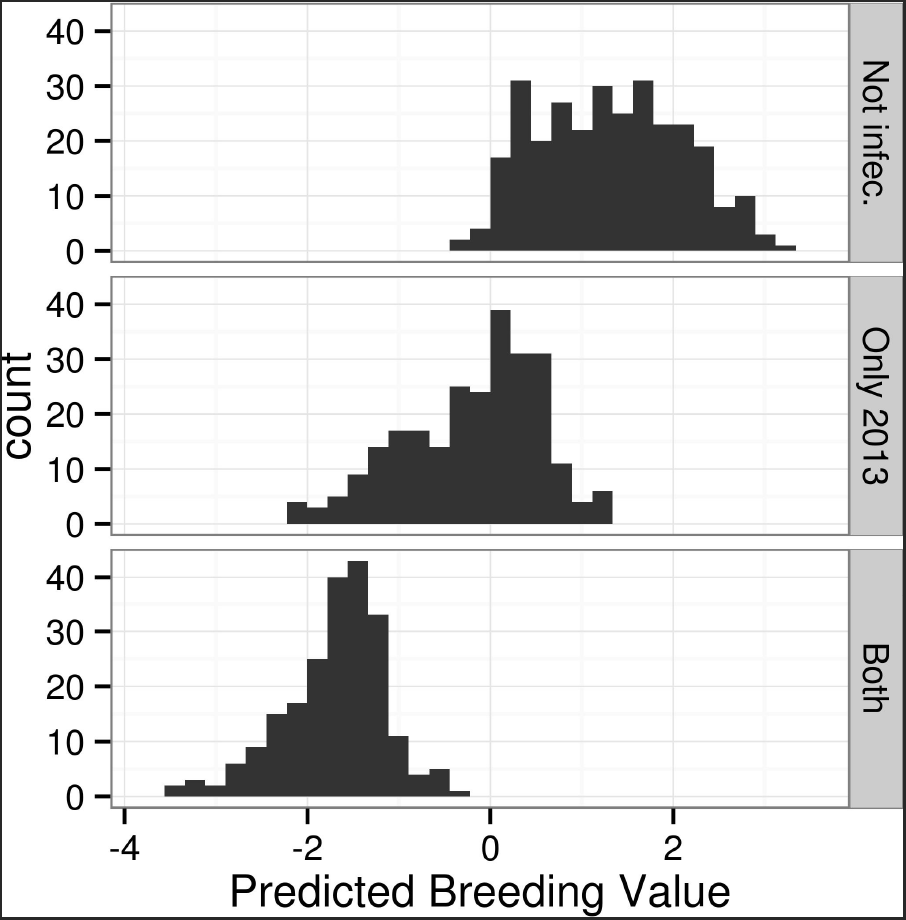
Distribution of predicted Individual Breeding Values (IBV) for the binomial component of Collar Lesions (CL, low values refer to high susceptibility), by observed infection status and history.

The ST component displayed very clear trends (Figure 10), particularly for the binary component, with the bottom rows showing a higher-than- average probability of remaining uninfected in both 2012 and 2013. Similarly, once infected, trees in the top rows were likely to display higher CL values than average. The patterns for both years were very similar, in accordance with the high mean posterior interannual correlation estimates: 0.82 and 0.89 for the binary and continuous components respectively.

**Figure 10:**
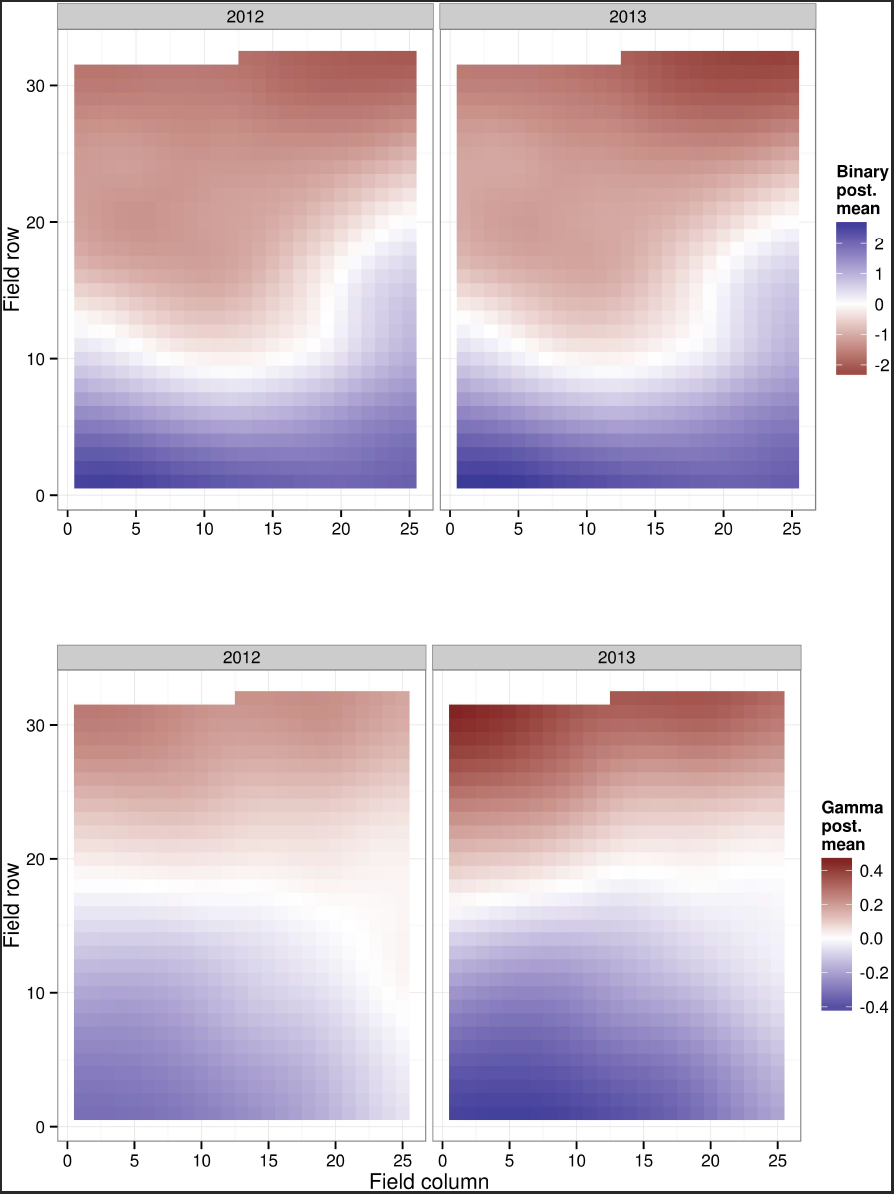
Posterior mean spatio-temporal (ST) Gaussian fields for the binary component (top) and the continuous component (bottom) of the model for Collar Lesions (CL), in the latent scale. Low values correspond to low susceptibility.

**Figure 7:**
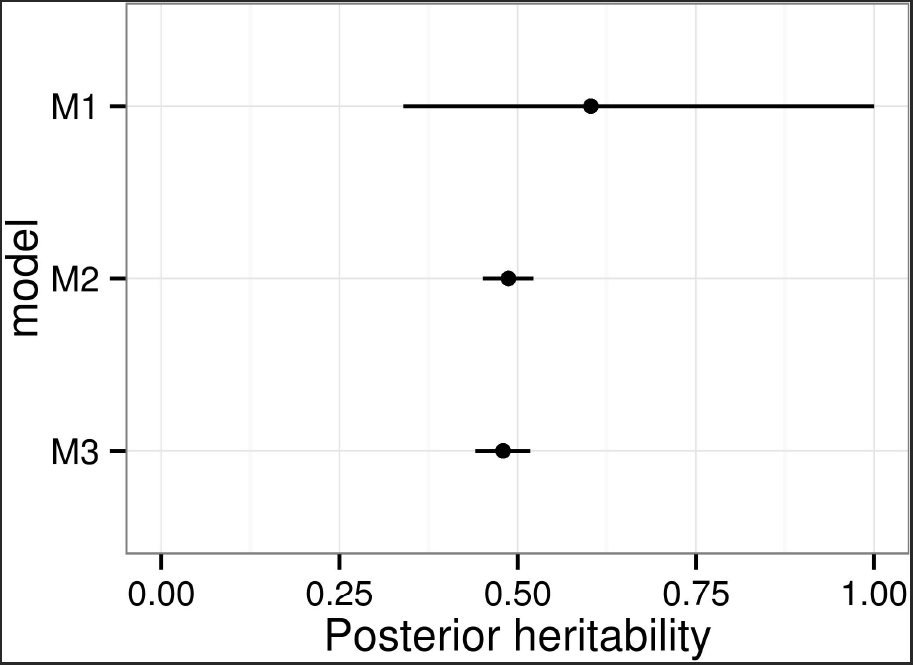
Posterior modes and 95% HPD Credible Intervals of the variance components for Crown Dieback (transformed: TCD).

In contrast to the model for TCD, the prior distribution played a more significant role in the predicted ST effect, particularly for the binary component (Figure 15, in the Annex). This was expected, as spatially-distributed binary data are very weakly informative.

**Figure 15:**
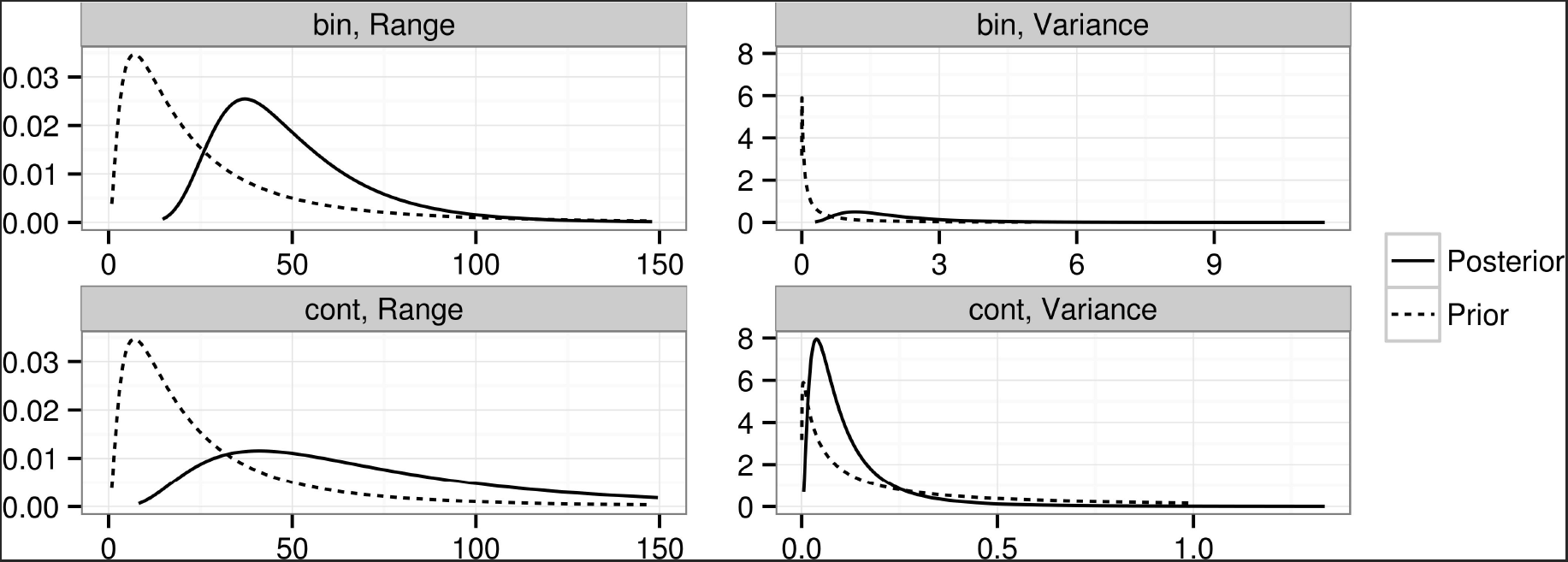
Prior and posterior densities for the Spatio-Temporal (ST) field characteristics of the binomial (bin) and continuous (cont) components of Collar Lesions (CL). Range unit = spacing between trees.

The mean posterior narrow sense heritabilities (*h*^2^) of CL were 0.49 and 0.42 for the binary and continuous components respectively, with a slightly wider 95% Credible Interval for the binary (0.32 - 0.64) than for the continuous (0.30 - 0.57) components. Having excluded the ST variance from the denominator, the posterior mean and Credible Interval for the heritability of the binary component increased to 0.60 (0.48 - 0.71).

### Genetic correlation between both symptoms

A moderate negative correlation was observed at the genetic level between TCD and CL. The individuals with highest genetic merit for TCD (i.e. those who are predicted genetically better-suited to resist the disease symptom in the crown) tend to have higher probability of avoiding collar lesion (r = 0.30, 95% CI: 0.25 – 1), and lower predicted genetic sensitivity to it, in the case of infection (r = -0.40, 95% CI: -1 – -0.34; Figure 11).

**Figure 11:**
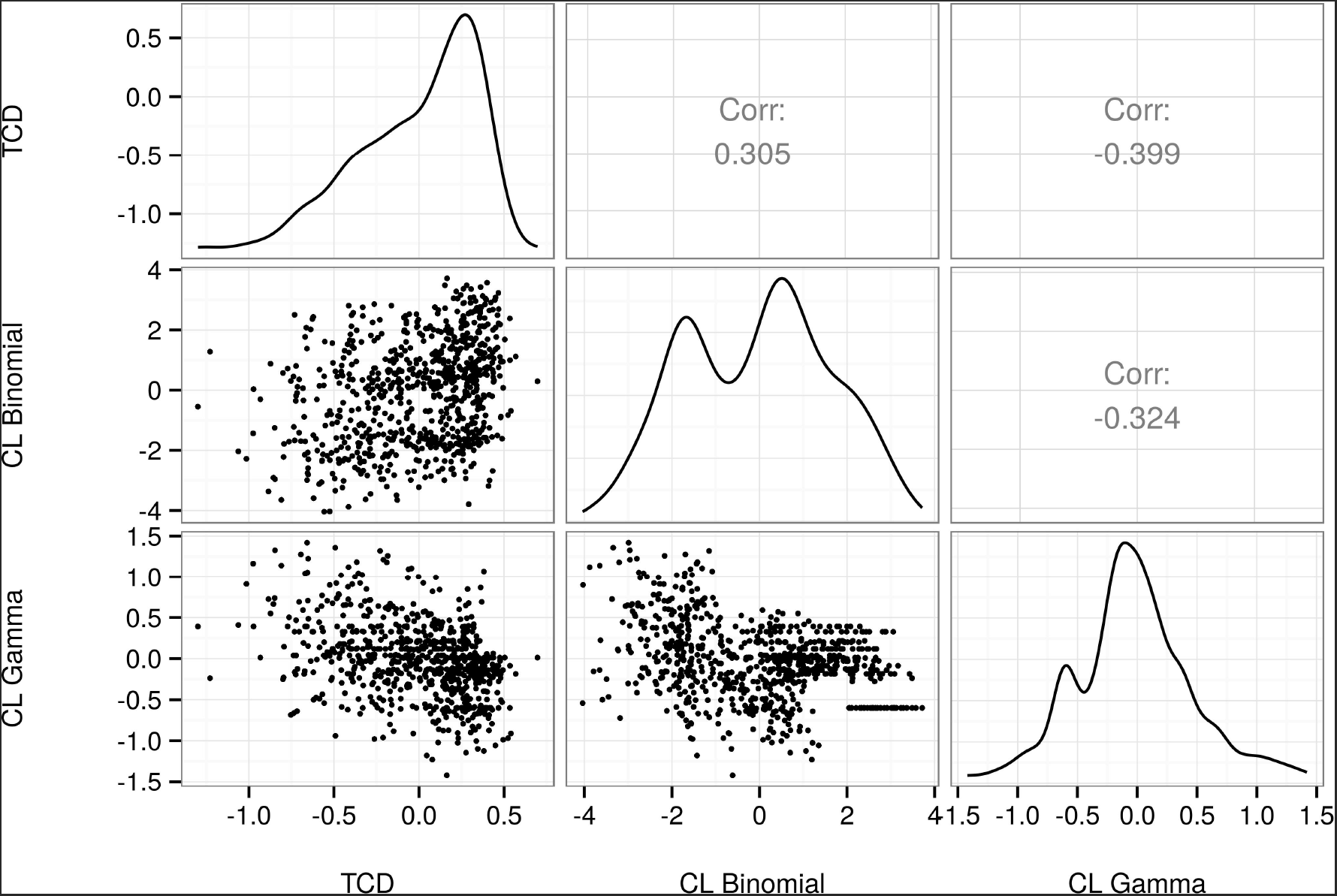
Correlation between individual breeding values (IBV) for Crown Dieback (transformed: TCD, low values refer to high susceptibility) and the binary (low values refer to high susceptibility) and continuous (low values refer to low susceptibility) components of Collar Lesions (CL).

## Discussion

### Disease progress and mortality

Only 3% of the trees died four years after the disease was detected in the experiment. Mortality rates ranging from 2% to 70% have been reported elsewhere (Enderle et al. 2013; Lobo et al. 2014; McKinney et al. 2014; McKinney et al. 2011; Metzler et al. 2012; Pliura et al. 2011; Pliura et al. 2014; Stener 2013), but very few studies allow to relate this rate to the time of exposure to the disease. Only three previous studies allow for comparisons, either because the trees started to be monitored for the disease right after planting or because initial proportions of symptomless tree suggest that the epidemics started recently. Two of these studies reported higher mortality rates despite similar times of exposure to the disease (9% in Enderle et al. 2013, 10% in Pliūra et al. 2011). However, they both analyzed younger trees (10 and 8 y.o., respectively) and it is commonly accepted that there is a negative correlation between susceptibility to *H. fraxineus* and tree age (Skovsgaard et al. 2010). By contrast, another study (Pliūra et al. 2014) reported a lower rate of 2% on very young trees (3 y.o.), but it was measured only one year after planting and on grafted material selected for resistance to the disease.

Looking at CD only yields a proportion of symptomless trees of 6% in 2014. However, already in 2013, more than 50% of the trees without visible CD had collar lesions. This result is in agreement with observations from Enderle et al. (2013) who found that 15% of otherwise healthy trees were affected by collar lesions. Removing trees without visible CD in 2014 but on which collar lesions were found the year before leads to a rate of symptomless trees of 4% only. Proportions of symptomless trees reported in the literature range from 1% (Lobo et al. 2014) to 58% (Kirisits et al. 2012). Again, comparisons should be made on trees of the same age that underwent similar exposure to the disease and on which monitoring of the disease was not restricted to the crown.

Two studies at least have reported health improvement on some trees (Lobo et al. 2014; Stener 2013). Indeed, a few percent of negative values were found for inter-annual increments for both CD and CL in the present study. A recovery process clearly acts for CD, when secondary and epicormics shoots are produced in reaction to the disease. By contrast, apparent remission in terms of CL is most certainly artifactual. Indeed, at early stages, collar lesions are often found below ground level and can be missed very easily.

### Collar lesions: still a lot to understand

In 2013, high proportions of trees showing only one of both symptoms demonstrate that they can occur independently. Nevertheless, a trend for a positive relationship between CD intensity and CL prevalenee was observed both in 2012 and in 2013. This result is consistent with previous findings (Bakys et al. 2011; Enderle et al. 2013; Skovsgaard et al. 2010).

Severities of both traits were also significantly positively correlated. Correlation at the phenotypic level has already been demonstrated by Bakys et al. (2011) who reported very high Pearson correlation coefficients (>0.57) between CD and several quantitative measurements of CL, and also by Husson et al. (2012) who reported Spearman correlation coefficients ranging from -0.2 to 0.7 in 60 natural plots. These last authors reported a tendency for higher correlation coefficients to happen in field plots with high overall mean CL. Using PCR assays, they also investigated the possibility that collar and branch lesions may be connected and did not find any evidence to support this hypothesis. Instead, they concluded on separate infection pathways with ascospores potentially infecting the stem base via lenticels in the bark.

There is still debate about whether *H. fraxineus* is the primary causal agent of collar lesions or not. Among other candidates, *Armillaria sp.* has been shown to occur at high frequency in ash collar lesions (Bakys et al. 2011; Husson et al. 2012; Lygis et al. 2005; Skovsgaard et al. 2010).

Although histological investigations would be needed, some results from the present study support the scenario where *H. fraxineus* induces the lesion while other fungi are responsible for secondary decay. Indeed, a significant positive genetic correlation between CD and CL suggests some common genetic determinism in the response of the host. Moreover, and as expected if *H. fraxineus* induced the disease, the genetic correlation coefficient was higher between CD and the continuous component of CL than with its binary component. Verification of this statement would benefit from measuring each lesion individually instead of expressing CL as a global girdling index. Of course, the possibility that resistance to *H. fraxineus* may also correlate with resistance to other collar-necrosis-inducing fungi cannot be excluded, and genetic variation for resistance to *Armillaria* has been demonstrated in at least one tree species (Zas et al. 2007).

On the other hand, the fact that spatial correlation happens at much shorter range for CD than for both components of CL can be seen as consistent with conclusions from Enderle et al. (2013). These authors stated that spatial autocorrelation for CL support the idea that collar lesions are caused by *Armillaria spp.* which spreads via the soil or via diseased roots while *H. fraxineus* would act as a more mobile secondary colonizer. Involvement of Armillaria could even explain why the ST Gaussian field revealed higher susceptibility in the upper part of the field experiment for CL in the present study whereas it was the reverse for CD, at least from 2010 to 2013. The upper part of the field experiment is, indeed, adjacent to a forested land which can be considered a reservoir for *Armillaria* whereas the bottom part is close to an agricultural land.

Other environmental factors, such as soil moisture which is known to correlate with CL (Husson et al. 2012), can also be accounted for by the ST Gaussian field. Moreover, they can influence the spatial distribution of BC, which is also autocorrelated in this field experiment (data not shown). This confounding factor would explain why despite the observed correlation between BC and CL prevalence, the inclusion of the variable in the model improves the DIC and WAIC while at the same time worsens the Deviance.

### Genetic components: their magnitude and how to estimate them properly

Only three studies explored the genetic variability of *F. excelsior* using OP progenies before this one (Kjær et al. 2012; Lobo et al. 2014; Pliūra et al. 2011). Among them, only one (Pliūra et al. 2011) found significant Provenance effects. This outlying result may be due to the fact that these authors studied a large number of provenances (24) covering a large distribution range across Europe. However, as stated by the authors, the observed Provenance effect may simply originate from the fact that local (Lithuanian) provenances had undergone natural selection by the pathogen before mother trees were selected whereas other European provenances did not. Other studies, including this one, involved fewer provenances, all of local origins, and did not conclude on significant Provenance effects.

All three previous studies concluded on significant Family effects and this is confirmed here. However, and because the methodology involves both familial (= mother tree) and individual (= tree) breeding values, the present study allows to conclude that most of the genetic variation lies within families.

The present study concludes on an *h*^2^ estimate for CD of 0.42 with a 95% HPD Credible Interval (0.38—0.47). Previous studies reported *h*^2^ estimates for CD in the range 0.20 – 0.49, excluding the 0.92 estimate provided by Pliūra et al. (2011) which was based on a global health assessment and was apparently overestimated due to early frosts. For CL, *h*^2^ estimates were 0.49 (binary component) and 0.42 (continuous component) with 95% Credible Intervals of about 0.3 – 0.6. These, however, cannot be compared to any data from the literature as this is the first time genetic variance components are estimated for this trait.

In the sake of accuracy, it is important to point out that the heritability estimates provided here rely on the assumption that the studied families are composed of half-sibs only. They could be biased upwards should the average relatedness within progenies be higher. Checking this hypothesis would require intensive genotyping, and to our knowledge only Kjær et al. (2012) conducted such verification.

Interestingly, the methodology implemented here allowed generating *h*^2^ estimates with fairly narrow Credible Intervals. We believe such procedure (i.e. explicit ST modeling under a Bayesian framework) should be used whenever possible, especially when infection started recently, meaning that (i) measured traits are not normally distributed due to a high proportion of asymptomatic or low infected trees and (ii) disease escape can be confounded with resistance.

Although moderate, these heritability estimates are much higher than those reported in *Ulmus* for resistance to *Ophiostoma novo-ulmi* (i.e. 0.14 +/- 0.06, Solla et al. 2015), especially when considering that *h*^2^ was estimated at the intra-specific level here whereas the figures in *Ulmus* come from a mating design involving not only the European *U. minor* species but also the Central-Asian species *U. pumila.* Although DED is a true vascular disease whereas *H. fraxineus* is not, comparison between both pathosystems seems relevant because in both cases the tree host species is facing a previously unknown fungal pathogen

Another relationship that was only partly explored here is the one between phenology and health status. In the case of *Ulmus*, percentage of wilting was negatively correlated with early bud flushing (Solla et al. 2015). In the case of *F. excelsior*, there are several indications that health status also correlates positively with early bud flushing but also with early leaf senescence and leaf coloring in the autumn (Bakys et al. 2013; McKinney et al. 2011; Pliūra and Baliuckas 2007; Stener 2013). A positive relationship was found between early BF and health status (CD) in the present study also. Although inoculation assays conducted by McKinney et al. (2012a) and Lobo et al. (2015) provided evidence that heritable *stricto sensu* resistance mechanisms occur in *F. excelsior*, some of the variability observed in the present study may thus come from phenological features leading to what should maybe termed ‘disease avoidance’. Although such features may prove valuable for selection, correlations between phenology and health status require further investigations as the disease can, in turn, modify the phenology of the host.

### Consequences for management and breeding

The present study adds to the common perception that no complete resistance to *H. fraxineus* can be found in *F. excelsior* but that there is significant variability for partial resistance and that it is heritable. Whether partial resistance is preferable to complete resistance will not be discussed here, but partial resistance is not always a guarantee of durability (Dowkiw et al. 2010). Most importantly, confirming that there is heritable genetic variation for susceptibility in *F. excelsior* is a prerequisite to save the species without introgressing resistance genes from exotic species like *F. mandshurica*, the supposed co-evolved host species of *H. fraxineus.*

Regarding natural regeneration, the positive consequence of moderate *h*^2^ estimates is that resistant *F. excelsior* trees that will remain or that will be reintroduced in natural populations will transmit resistance to the next generation in a highly additive manner. Of course, highly susceptible trees will also transmit their behavior to the next generation. However, within family variation is so high that removing highly susceptible adult trees does not seem to be necessary except to avoid falling hazard. Promoting regeneration in natural stands and letting natural selection to occur may prove sufficient. Nevertheless, since selection by the pathogen will occur at the seedling stage, this optimistic scenario may not come true if juvenile-adult correlations for resistance are low. These correlations have not been evaluated at the individual level yet, but juvenile trees are known to be more susceptible to the disease (Skovsgaard et al. 2010).

For planting, high levels of intra-familial variation suggest that seed material from selected parents planted in seed orchards would require further selection. This could be done either in the nursery or directly in the field if planting is done at higher density than usual. Although not common for Ash (except for ornamental trees), clonal selection would certainly allow higher genetic gain. However, several questions would then arise. First is that of the number of clones to release to ensure sufficient genetic variability, not only to avoid resistance breakdown but also to cope with future threats like the Emerald Ash Borer (*Agrilus planipenis*) and to combine Ash dieback resistance with other traits of interest. Second is that of the techniques to use to propagate the selected Ash clones and the associated costs. *F. excelsior* is much easier to propagate through grafting than through cuttings. Moderate genetic correlations between CD and CL and moderate heritabilities for both traits suggest having two separate selection schemes, one for rootstocks (CL) and one for grafts (CD). Ash propagation through cuttings would be less expensive and certainly deserves more investigation should clonal selection be considered.

It is important to consider also that all genetic variance estimates were measured at a given point in time and in a given environment. *H. fraxineus* having a sexual stage, new variants of the pathogen appear each year and thus adaptation in the pathogen’s populations can occur. This may explain why significant GxE can be observed for resistance to *H. fraxineus* and why cloned (grafted) resistant material can become susceptible after only two growing seasons (Pliūra et al. 2014).

## Acknowledgements

This study was essentially supported by the French Ministry of Agriculture (Programme 149 – Action 13 – Sous action 32). F. Muñoz is partially funded by research grant MTM2013-42323-P from the Spanish Ministry of Economy and Competitiveness and ACOMP/2015/202 from Generalitat Valenciana (Spain). We thank the town of Devecey for providing the land to install the field experiment and the technical team of INRA Nancy for their continuous commitment.

## ANNEX: Model comparisons and diagnostic plots

### Models for Crown Dieback

Model M1 was used as a reference model as it uses only unstructured random effects, which is the most basic setting. It includes fixed Year and Block effects and unstructured random Family and Family X Block effects. Initially, we also included fixed Provenance and Provenance x Block effects but the Provenance effect was not significant and a Likelihood Ratio Test confirmed that this model was not significantly better than M1 (χ^2^(22) = 31.176, *p* = 0.093).

Model M1 was also fitted using REML (Bates et al. 2015), to compare results with previous literature using the same methodology. We checked that Bayesian and frequentist point estimates of the variance components were similar and thus we report only the Bayesian results.

In model M2, the Family x Block interaction effect was also removed since its effect was completely absorbed by the individual genetic effect.

Model M3 replaced the unstructured Block effect by a Spatio-Temporal random effect.

Models M4 and M5 included, in addition, two potentially explanatory variables. Namely, the Provenance and the Bud-Flush precocity, which entered as fixed effects, respectively.

Lastly, Model 6 forces an explicit null Family effect by imposing a sum-to-zero constraint in the breeding values for each family. Comparing this model with M5 allows testing the hypothesis of significant differences in mean genetic values between families.

The results from model M1 were not reliable since the diagnostics on the residuals were invalidating. In particular, a Shapiro-Wilk normality test yielded a *p*-value of 2.2e — 16, which is in the limit of the floating-point number representation. The high posterior uncertainty on the narrow-sense heritability (*h*^2^) was very high, the only conclusion being that the heritability is likely to be above 0.30. Consistently, a frequentist REML inference on this model yielded a bootstrap estimate of *h*^2^ of 0.61, and a 95% Confidence Interval of 0.23–0.98.

Model M2 represented a huge improvement in the three comparison criteria and led to a much narrower Credible Interval for ñ^2^. But model M3, including the ST effect, was significantly better.

Model M4 yielded worse values of all the comparison criteria, suggesting that there is no Provenance effect.

Model M5 improved the DIC and the WAIC, but yielded a higher Deviance. Since there is prior evidence in the literature of a relationship between bud-flush precocity and CD (Bakys et al. 2013; McKinney et al. 2011; Pliura and Baliuckas 2007; Stener 2013), we decided to include this variable in the model.

M6 overfits the data as it turns out from the almost double effective number of parameters, which is comparable in magnitude with the total number of observations. Under these conditions the DIC and WAIC criteria are not reliable (Plummer 2008). The marginal likelihood being considerably smaller than that of model M5, the variance between families cannot be considered to be zero.

### Models for Collar Lessions

To check the relevance of the Basal Circumference (BC) as an explanatory variable, different parameterizations of this factor were used. First (model M1.1), BC was split into 6 categories (≤ 30,]30,45], ]45,60], ]60,75], ]75,90], >90); model M1.2 also considered also the interaction with the Year; models M1.3 and M1.4 used BC as linear and quadratic regressors respectively; finally, model M1.5 considered a non-parametric random function of the variable using a second-order random walk.

The relevance of including a fixed Provenance effect was tested by including either its main effect alone (model M2.1) or its main effect plus its interaction with the Year (model M2.2).

